# CrebA regulation of secretory capacity: Genome-wide transcription profiling coupled with in vivo DNA binding studies

**DOI:** 10.1101/2025.06.12.659381

**Authors:** D.J. Jackson, D. Peng, S.A. Shinde, A. Holenarasipura, P. Cahan, D.J. Andrew

## Abstract

DNA binding assays, expression analyses, and binding site mutagenesis revealed that the Drosophila CrebA transcription factor (TF) boosts secretory capacity in the embryonic salivary gland (SG) through direct regulation of secretory pathway component genes (SPCGs). The mammalian orthologues of CrebA, the Creb3L-family of leucine zipper TFs, not only activate SPCG expression in a variety of mammalian tissues but can also activate SPCG expression in Drosophila embryos, suggesting a highly conserved role for this family of proteins in boosting secretory capacity. However, in vivo assays reveal that CrebA binds far more genes than it regulates, and it remains unclear what distinguishes functional binding. It is also unclear if CrebA is the major factor driving SPCG gene expression in all Drosophila embryonic tissues and/or if CrebA also regulates other tissue-specific functions. Thus, we did single cell RNA sequencing (scRNA-seq) of wild-type (WT) and *CrebA* null embryos to explore the relationship between CrebA binding and gene regulation. We find that CrebA binds the proximal promoters of its targets, that SPCGs are the major class of genes regulated by CrebA across tissues, and that CrebA is sufficient to activate SPCG expression even in cells that do not normally express the protein. A comparison of scRNA-Seq to other methods for capturing regulated transcripts reveals that the different methodologies identify overlapping but distinct sets of CrebA targets.

## Introduction

Protein secretion is essential for maintenance of tissues and homeostasis of organisms. Whereas all cells secrete proteins, some secrete extraordinary amounts as constituents of dedicated secretory organs. For example, the human pancreas produces both the hormones that regulate circulating levels of blood sugar and the large quantities of the enzymes necessary to digest food (Karpinska and czauderna 2022). The mammary glands are specifically designed to produce high quantities of milk in a highly regulated fashion (Peaker 2002). These dedicated secretory organs require unique adjustments to common cellular processes to accommodate the high-level synthesis of secreted products. This requires coordination among the transcription factors (TFs) that regulate expression of the machinery of secretion and of translation, as well as the factors that determine what kind and how much secretory product will be made. To date, the TFs mediating secretory function of only a very few organs have been discovered (Abrams and Andrew 2005; Fox *et al*. 2010; Barbosa *et al*. 2013; Khetchoumian *et al*. 2019; Wang *et al*. 2024) and an understanding of how these factors coordinate their activities during the early stages of development to meet the immediate and long-term physiological demands of each organ is only beginning to be revealed.

The Drosophila salivary gland (SG) is a premier example of a simple dedicated secretory organ that requires high levels of coordinated regulation throughout development to properly carry out its secretory function (Loganathan *et al*. 2021). Studies of the SG over the past few decades have revealed key transcription factors (TFs) controlling both secretory capacity and secretory specificity (Abrams and Andrew 2005; Fox *et al*. 2010; Fox *et al*. 2013; Johnson *et al*. 2020; Loganathan *et al*. 2022). The CrebA leucine zipper TF boosts secretory capacity in the SG as well as in other professional secretory cells of the Drosophila embryo (Abrams and Andrew 2005; Fox *et al*. 2010). Loss of *CrebA* results in a reduction of SG lumen volume, structural defects in the cuticle secreted by epidermal cells, and significant decreases in expression of secretory pathway component genes (SPCGs) (Abrams and Andrew 2005; Fox *et al*. 2010; Johnson *et al*. 2020). The regulatory role of this family of TFs is also conserved in vertebrate systems, as all five CrebA mammalian orthologs have been shown to regulate SPCGs in various tissues (Murakami *et al*. 2009; Saito *et al*. 2009; Barbosa *et al*. 2013; Reiling *et al*. 2013; Khetchoumian *et al*. 2019; Sampieri *et al*. 2019). Importantly, all five human CrebA orthologs also have the capacity to induce expression of Drosophila SPCGs in the Drosophila embryo (Fox *et al*. 2010; Barbosa *et al*. 2013), suggesting that their mechanisms of action are also conserved.

SPCGs encode the protein machinery of secretion (**Figure 1**), including components of the signal recognition particle (SRP), the SRP receptor (SR), the machinery for co-translational transport of nascent polypeptides into the endoplasmic reticulum (ER), the Cop-I and Cop-II machinery for retrograde and anterograde trafficking of secretory vesicles, and the proteins and enzymes that populate the ER and different Golgi compartments to provide organelle structure or to covalently modify secreted and transmembrane proteins. CrebA boosts secretory capacity through direct binding and transcriptional activation of SPCGs and of other genes, including for example Xbp1 (Johnson *et al*. 2020), a TF best known for its role in the unfolded protein response (UPR) (Glimcher *et al*. 2020) but that has also been implicated in boosting secretory capacity in exocrine glands (Lee *et al*. 2005). Microarray studies assessing overall changes in gene expression in *CrebA* null embryos also revealed that transcripts for many genes encoding secretory cargo are also considerably decreased with loss of *CrebA* (Fox *et al*. 2010). However, although binding of CrebA is essential for direct regulation of gene expression, binding and regulation of specific genes are not tightly correlated. A previous in situ analysis showed that a subset of SG-expressed genes that are directly bound by CrebA had no overt changes in their SG expression in the absence of CrebA (Johnson *et al*. 2020). Thus, whereas CrebA plays a major role in boosting secretory capacity in the SG, it remains unclear how many CrebA-bound genes are affected by loss of *CrebA* and if genes not bound by CrebA are also affected by *CrebA* loss.

**Figure 1.**
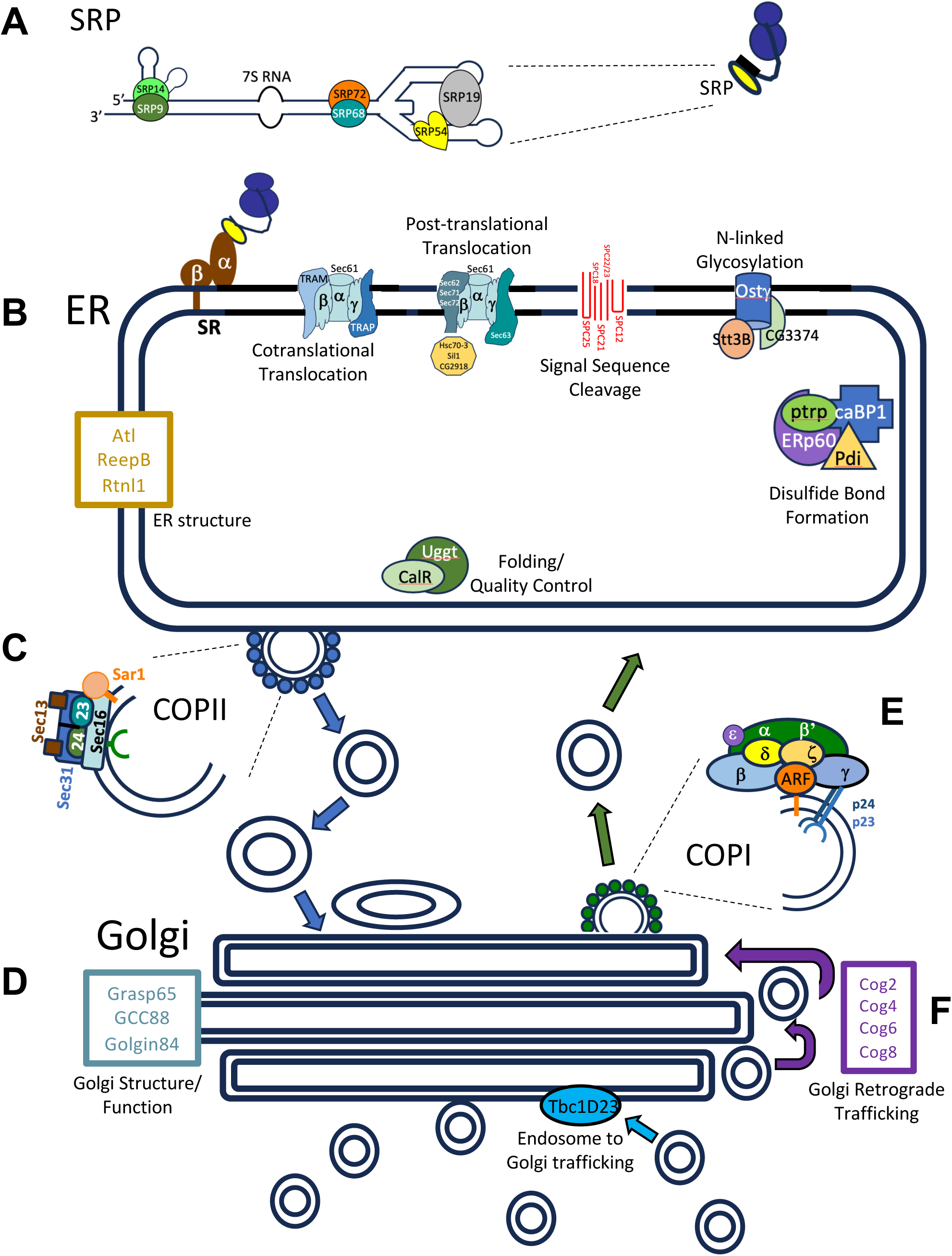
Cartoon of the secretory pathway showing the major protein complexes and individual proteins associated with each complex. **A.** The Signal Recognition Particle (SRP) is composed of several SRP proteins and a 7S RNA. As shown in the smaller cartoon to the right, SRP (yellow oval) binds the N-terminal hydrophobic signal sequences (black rectangle) of nascent polypeptides (black line) emerging from ribosomes (blue circles). **B.** The bound SRP complex subsequently docks at the SRP receptor (SR) on the endoplasmic reticulum (ER). The ribosome bound nascent polypeptide is then handed off to the machinery for cotranslational translocation (Sec61 complex and associated proteins), where the elongating polypeptide is threaded into the ER lumen as it is synthesized. Sec61 is also part of the machinery for post-translational ER translocation. As proteins are threaded into the ER lumen, chaperones assist in correct folding (yellow decagon). The N-terminal signal sequence is removed by the SPase complex (Red lines). Several post-translational modifications of proteins occur in the ER – shown modifications include N-linked glycosylation, disulfide bond formation, ER-folding, and quality control. Other proteins are important for overall ER structure – tubular or planar (Gold box). ER cargo proteins are carried to the Golgi through vesicular trafficking. **C.** Formation and release of ER vesicles destined for the Golgi (small blue circles surrounding two larger circles) is mediated by the Coat Protein Complex II (COPII). **D.** Proteins important for overall Golgi structure and function are shown in the blue/gray box. **E.** Retrograde vesicular trafficking from the Golgi back to the ER is mediated by the Coat Protein Complex I (COPI) (small green circles surrounding two larger circles). Retrograde vesicular trafficking in the Golgi is mediated by the Conserved Oligomeric Golgi complex (COG) proteins (purple box).

Here, we utilized our previously established scRNA-seq datasets from WT embryos (Peng *et al*. 2024) to examine *CrebA* expression across tissues and found that levels of CrebA closely correlated with relative levels of SPCG expression. We then carried out single cell RNA sequencing of *CrebA* null embryos during early organogenesis to identify tissue-specific changes in gene expression linked to loss of *CrebA*. These expression studies revealed that tissues with the highest relative SPCG expression in WT embryos showed the most drastic reduction in SPCG expression in *CrebA* mutants. This analysis also revealed unexpected differences in specific targets uncovered by this methodology versus previously used transcriptomic analyses. We find that CrebA can induce ectopic expression of SPCGs in tissues that do not normally express any detectable CrebA, but we observed tissue-, stage-, and gene-specific differences in SPCG activation driven by ectopic CrebA. Finally, we revisited previously generated CrebA ChIP-seq DNA binding data to identify motif features that correlate with CrebA binding and regulation versus CrebA binding without corresponding regulation. Our analyses reveal that CrebA activates gene expression through binding to promoter proximal regions that contain CrebA consensus sites defined by previous in vitro DNA binding assays. Our analyses of overall gene expression changes and DNA binding studies support a model wherein CrebA primarily functions as a transcriptional activator and that its major role across all tissues in which it is expressed is to boost secretory capacity.

## Materials and Methods

### Fly Stocks

Two loss-of-function alleles for *CrebA* were used in this study. The *CrebA^23w-^* allele was generated by excision of a P-element insertion 772 nt upstream of the transcription start site resulting in the deletion of the region flanking the 5’ end of the *CrebA* gene (Andrew *et al*. 1997). *CrebA^03576^* (RRID:BDSC_10183) is a 14.5 kb P-element insertion in the first intron of the CrebA gene (Rose *et al*. 1997). Both alleles are protein null based on immunostaining with CrebA antiserum (Andrew *et al*. 1997; Rose *et al*. 1997). Gal4 lines for ectopically expressing UAS-*CrebA* (Rose *et al*. 1997) were obtained from the Bloomington Stock Center: *en*-Gal4 (RRID:BDSC_1973), *Mef2*-Gal4 (RRID:BDSC_27390), *twi*-Gal4 (RRID:BDSC_914), elav-Gal4 (RRID:BDSC_458) and *nos*-Gal4 (RRID:BDSC_4442). UAS-CrebA (RRID:BDSC_32572) was generated by Sarah Smolik.

### Drosophila embryo collection

*CrebA^23w-^/TM3, twi-GFP* and *CrebA^R3576^/TM3, twi-GFP* were crossed to generate CrebA protein-null embryos for processing using the same single-cell RNA-sequencing pipeline previously established in the lab (Peng *et al*. 2024). Briefly, embryos were collected and aged to capture stages 10-12 of embryonic development (“early organogenesis”) or stages 13-16 (“late organogenesis”). Embryos were dechorionated in a 50% bleach solution and sorted using a COPAS Flow Pilot sorter to isolate GFP-negative (CrebA protein null) embryos. Live embryos were visually inspected under a fluorescent microscope to confirm both stage and absence of GFP-positive individuals. Samples were immediately prepared for single cell isolation and 10X Genomics single-cell RNA sequencing.

### Cell isolation

The cell isolation protocol was adapted using previously described methods (Karaiskos *et al*. 2017; Peng *et al*. 2024). Pooled embryos were resuspended in 1 mL of chilled PBS and dissociated using a loose pestle in a glass dounce homogenizer (Wheaton catalog #: 23ND77). Samples were centrifuged at 1500 RCF at 4°C for 3 minutes and resuspended in 1 mL of chilled PBS. Cell suspensions were gently passed through a 22G x 2” needle mounted on a 5 mL syringe 20 times followed by an additional four passes through a 23G x 2“needle to further dissociate cells. Cells were centrifuged for 3 min at 3300 RCF in a tabletop centrifuge and resuspended in 100 µL of chilled PBS. After resuspension, cells were filtered through a 40 µM strainer (Greiner 542140) into a 1.5 mL microcentrifuge tube. Samples were immediately transported on ice to the Johns Hopkins Single Cell & Transcriptomics Core where cell viability was assessed using a Countess 3 Cell Counter before processing for 10X Genomics single-cell RNA sequencing. At least two biological replicates were generated for each experimental condition representing early and late organogenesis for *CrebA* mutant embryos. Unfortunately, the late organogenesis samples were not of sufficiently high quality to analyze in detail. Thus, we focus on WT and early stage *CrebA* mutant data for which we obtained two high quality biological replicates in this report.

### Single cell RNA-seq processing

10X CellRanger software (Zheng *et al*. 2017) was used to align the raw scRNA sequence data to the Drosophila reference genome (v 6.33) from FlyBase (Gramates *et al*. 2022). Cells were pre-processed using R (v 4.1.2) and Seurat (v 4.2.1) before conducting a batch integration on each sample. Cells with high mitochondrial transcript expression (> 0.25) were classified as poor-quality and removed from subsequent analyses. A standard Seurat workflow was used to cluster cells above the average log2-fold change of 0.25 (due to potentially constituting empty droplets). DoubletFinder (Mcginnis *et al*. 2019) (version 2.0.3) was used to remove doublets and filter out any remaining low-quality cells. After completing quality control measures, Seurat datasets from biological replicates were merged and log-normalized using standard Seurat function for subsequent principal component analysis (PCA). Harmony (Korsunsky *et al*. 2019a) (version 0.1.0) was used to correct batch effects between biological replicates and corrected principal components were then used to construct UMAPs and Louvain clustering. Differential gene expression analysis between *CrebA* mutant and wild-type tissues were performed using Presto (v 1.0.0) (Korsunsky *et al*. 2019b). Gene set enrichment analysis was performed using fgsea package (v 1.20.0) using default parameters (Korotkevich *et al*. 2021). The gene set library “GO_Biological_Process_2018” was downloaded from modEnrichR (Kuleshov *et al*. 2019), and gene sets with fewer than 10 or more than 500 genes were not included in the analysis.

### Manual annotation of cell types

Clusters were manually annotated by using the Berkeley Drosophila Genome Project (BDGP) in situ hybridization data to assign cell type based on tissue-specific expression profiles of the top 20-50 differentially expressed genes from each cluster. Cell typing was further validated by cross-referencing the expression profile of each cluster with our previously established WT scRNA-seq database (Peng *et al*. 2024).

### Integration of ChIP-seq and scRNA-seq data

The processed narrowPeak files for fkh-Gal4>UAS-CrebA-GFP embryos with anti-GFP (fkh_CrebA_peaks) and sage-Gal4>UAS-CrebA-GFP embryos with anti-GFP (sage_CrebA_peaks) ChIP-seq dataset were downloaded from GEO (accession ID: GSE141778) (Johnson *et al*. 2020). The overlapping peaks between “sage_CrebA_peaks” and “fkh_CrebA_peaks” were found using bedtools (v2.31.1) (Quinlan and Hall 2010) (keep original entry for “sage_CrebA_peaks”) as salivary gland specific CrebA binding peaks. Genes with 5’ transcription starting site (TSS) closest to the center of the intersecting peaks (both upstream and downstream and less than 1 kb away) were assigned as CrebA bound genes in salivary gland. Among the pool of CrebA bound genes, genes (compared between *CrebA* mutant and wild-type salivary gland cells) with logFC < -0.15 and p-val < 0.05 were identified as activated genes and genes with logFC > 0.15 and p-val < 0.05 were identified as repressed genes in scRNA-seq. In addition to activated genes found using scRNA-seq, genes that were previously identified to be activated by CrebA via in situ in salivary gland cells and microarray were also included (Abrams and Andrew 2005; Fox *et al*. 2010; Johnson *et al*. 2020). To ensure the activated genes from whole-embryo microarray are regulated in salivary gland cells, the microarray-identified CrebA activated genes (fold-change < -1.25 and p-val < 0.05) were further filtered down to those that are expressed in at least 10% of the salivary gland cells in the wild-type scRNA-seq data and show a decrease in expression (logFC < 0) in the mutant salivary gland cells in the *CrebA* mutant scRNA-seq.

Gene set enrichment of CrebA bound-activated and bound-repressed genes was performed using enrichR (v 3.1) (Kuleshov *et al*. 2016). DREME (v 5.5.4) (Bailey 2011) was used to identify motifs in CrebA binding regions associated with bound and activated genes, bound and repressed genes, and bound and no change genes. Prior to running DREME, the CrebA binding regions were trimmed to be -250 and +250 base pairs around the center of the intersecting peaks. The no change genes were defined as 150 least differentially expressed (smallest logFC) genes between mutant and wild-type salivary gland cells. TOMTOM (v5.5.4) (Gupta *et al*. 2007) was used to compare motifs against a database of transcription factors (Fly Factor Survey) with known motifs (Zhu *et al*. 2011).

### In situ hybridization experiments

In situ hybridization was done according to the methods used by the Berkeley Drosophila Genome Project (BDGP), with technical advice provided by R. Weiszmann (Weiszmann *et al*. 2009). Antisense RNA probes were generated as described in (Lehmann and Tautz 1994) from cDNA clones obtained from the Drosophila Genomics Resource Center (DGRC). cDNA clones were sequenced prior to use to confirm their identity and included cDNAs encoding the following SPCGs: *SrpRα* (LD25651), *Sec61β* (RH61539), *Spase12* (RE02772), *KDEL-R* (LD06574), and *βCop* (GH09317).

## Results

### Expression of CrebA correlates with SPCG expression in WT stage 10-12 embryos

Single-cell RNA sequencing (scRNA-seq) was previously applied to whole wild-type Drosophila embryos to establish a comprehensive single-cell atlas from when organogenesis commences through organ maturation (Peng *et al*. 2024). Summarized, wild-type embryos representing early (stage 10-12) and late (stage 13 – 16) organogenesis were collected and broken down to single-cell resolution for scRNA-seq. Through this analysis, 35 clusters corresponding to 23 broad cell types were identified in early samples (**Figure 2A**) and 36 clusters corresponding to 29 broad cell types were identified in late samples (**Figure S1A**). A comparison of *CrebA* transcript levels during early organogenesis revealed that tissues with the highest expression of *CrebA* are the salivary gland (SG), amnioserosa, plasmatocytes, fat body, trachea, glia, and hindgut (**Figure 2B**). Very low levels are detected in most other tissues with no expression in germ cells, crystal cells, circular visceral mesoderm (cVM), gut endoderm, central nervous system (CNS), and somatic muscles. The expression data are largely consistent with *CrebA-*expressing cell types identified based on in situ hybridization (**Figure 2C**, left panels), except that glial and hindgut expression are not visually obvious at these stages. However, *CrebA* mRNA in hindgut and glial cells can be detected during stages 13-16 (**Figure S1B**). Correspondingly, stage 13-16 embryo scRNA-seq data indicates that SG, plasmatocytes, fat body, trachea, glia, and hindgut cells continue to express high levels of *CrebA*, whereas a decrease in expression of *CrebA* is observed in amnioserosa (**Figure 2B, S1B**), correlating with the contraction and eventual death of this tissue during these stages (Lacy and Hutson 2016). New tissues that were not distinguishable at early stages also express relatively high levels of *CrebA* during stages 13 – 16, including the epipharynx, hypopharynx and esophagus. Staining of embryos with CrebA-specific antiserum (Fox *et al*. 2010) reveals that the pattern of CrebA protein accumulation parallels that of *CrebA* transcripts both early and late (**Figure 2C, S1C**). Thus, *CrebA* regulation during embryogenesis is at the transcriptional level.

**Figure 2.**
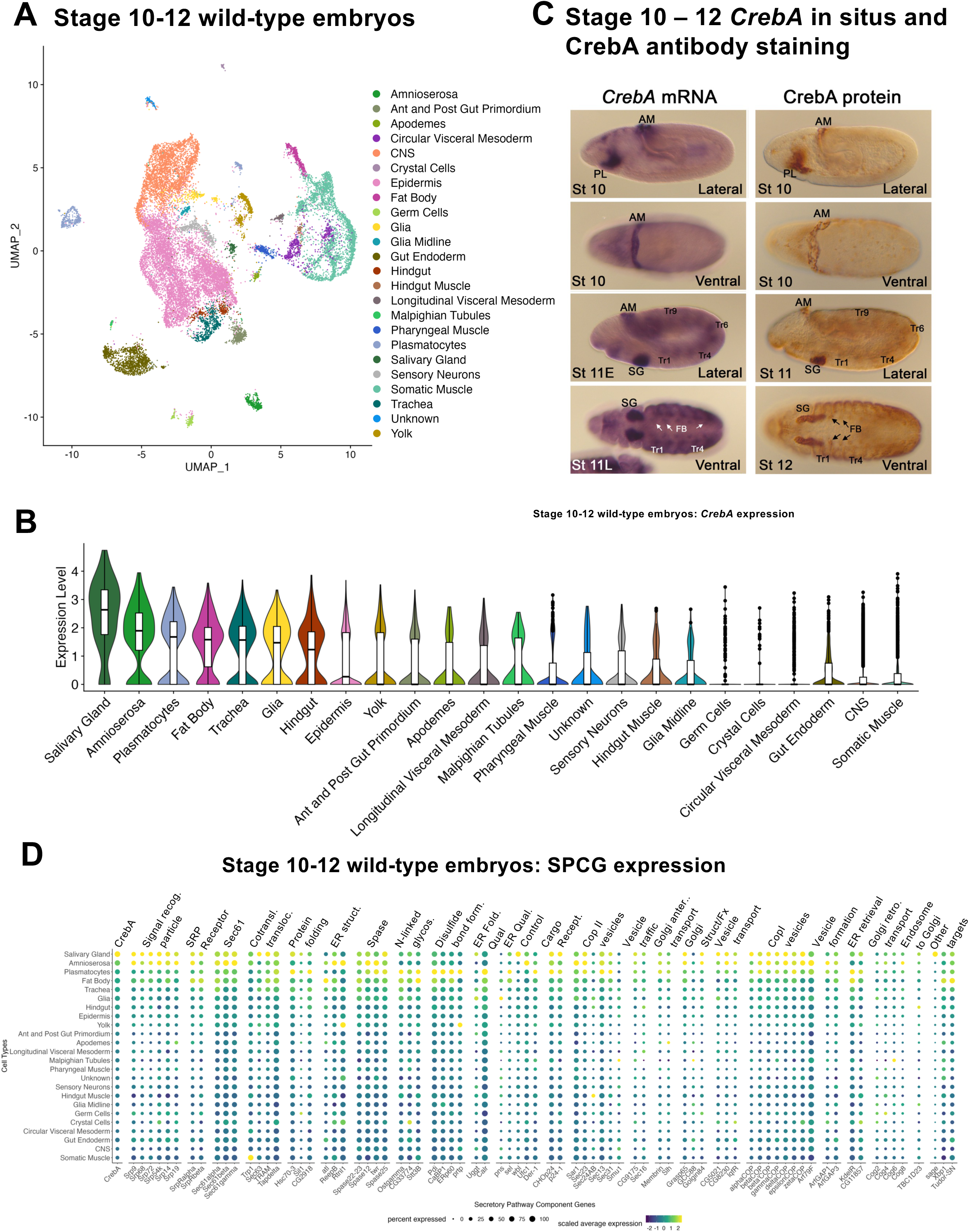
Analysis of CrebA and SPCG expression across tissues from early-stage (stage 10-12) wild-type embryos. **A.** UMAP showing that all the major cell types are represented in the wild-type stage 10 – 12 data (Peng et al., 2024). **B.** Violin plot of CrebA expression across tissues: tissues expressing the highest levels are on the left, lowest on the right. **C.** In situ hybridization of *CrebA* (left panels) and CrebA immunostaining (right panels) at the stages captured by the early scRNA-seq analysis. The first and third rows are lateral views, the second and fourth are dorsal-ventral views. **D.** Early embryonic expression levels of *CrebA*, most known SPCGs, and three other known CrebA targets across early embryonic tissues. Only genes that are sufficiently expressed in at least one cell type are shown. Abbreviations: AM – amnioserosa, PL – plasmatocytes, SG – salivary gland, Tr – trachea, FB – fat body, St – stage.

Since CrebA is known to boost expression of secretory pathway component genes (SPCGs) (Abrams and Andrew 2005; Fox *et al*. 2010; Johnson *et al*. 2020), we next examined the relative tissue-specific expression levels of previously identified SPCGs as well as other known CrebA targets (*sage*, *Xbp1*, and *Tudor-SN*) in the early WT single-cell atlas. As expected, our scRNA-seq data revealed that the SG-specific TF gene *sage* is exclusively expressed in the SG cluster (**Figure 2D, S1D**), consistent with both in situ detection of the *sage* transcript and immunostaining for the Sage protein (Abrams *et al*. 2006; Fox *et al*. 2013). *Xbp1* and *Tudor-SN* shared similar tissue levels of expression with *CrebA*, with highest expression seen in the SG, amnioserosa, plasmatocytes, and fat body (**Figure 2D**). Importantly, similar trends were also seen across the various SPCGs; highest levels of expression were observed in the SG, amnioserosa, plasmatocytes, fat body, trachea, and glia, and relatively low-level expression in other tissues, with some rare expression of a small subset of these genes in one or more tissues (**Figure 2D**). Similar correlations between expression of *CrebA* and SPCGs were also observed at later stages (**Figure S1D**); tissues with high levels of SPCG expression also had high levels of CrebA. A single exception was the garland cells, which express very high levels of SPCGs but lower levels of CrebA (**Figure S1D**). Notably, garland cells express the highest level of *Xbp1* (**Figure S1D**), a related bZip TF implicated in SPCG upregulation in mammalian exocrine glands and in zebrafish cement glands (Lee *et al*. 2005; Wang *et al*. 2024). Altogether, this analysis suggests that CrebA could upregulate SPCGs in all the tissues in which it is expressed.

### Single-cell RNA-sequencing in *CrebA* mutant embryos

Next, we explored the transcriptional consequences when CrebA is absent in the embryo. To generate scRNA sequencing data from *CrebA* null embryos, adults carrying either of two unrelated protein null alleles of CrebA (**Figure 3A**) in trans to a GFP-tagged balancer chromosome were crossed to obtain *CrebA* null embryos (*CrebA^23w-/03576^*) for processing through our previously established scRNA sequencing pipeline (Peng *et al*. 2024). Embryos were collected, dechorionated and processed through a COPAS Flow Pilot sorter to isolate *CrebA* null GFP-negative individuals. After visual inspection under a fluorescent microscope to confirm successful sorting and appropriate staging, live samples were prepared for single cell isolation and 10X Genomics single-cell RNA sequencing. After the removal of doublets and low-quality cells via standard scRNA-seq processing measures (see Materials and Methods), 24,523 cells (Median UMI count: 7042) were recovered from two biological samples of stage 10-12 *CrebA* mutant embryos (**Figure S2A**). In total, we identified 23 distinct cell types (**Figure 3B**) from 40 cell clusters (**Figure S2B**) through matching the top differentially expressed (DE) genes from each cluster (**Table S1**) to in situ hybridization data from Berkeley Drosophila Genome Project and to our previously established *Drosophila* embryos atlas dataset.

**Figure 3.**
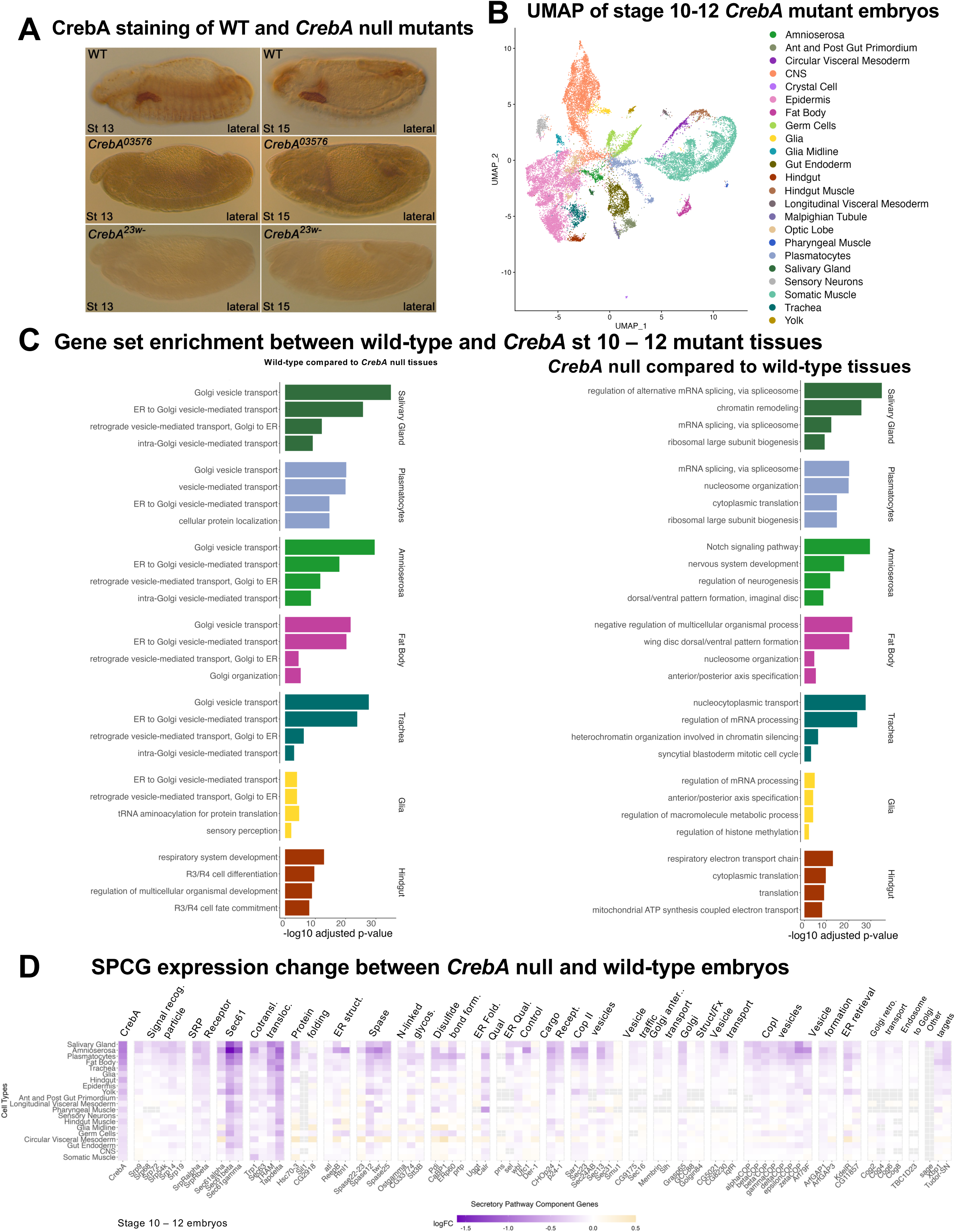
Analysis of gene expression changes in *CrebA* null versus wild-type early embryos (stage 10-12). **A.** CrebA immunostaining of late-stage WT (top panels) and *CrebA* null mutant embryos (middle and lower panels). **B.** UMAP from early stage *CrebA* null embryonic scRNA-seq showing the major cell types represented in the data. **C.** Gene set enrichment analysis (GSEA) comparing genes expressed in individual tissues from WT and CrebA null early embryos. The left bar plots correspond to biological processes enriched in the WT relative to *CrebA* null tissues; the right bar plots correspond to biological processes enriched in *CrebA* null tissues relative to WT. **D.** Relative changes in expression of *CrebA*, most known SPCGs, and three other known CrebA targets across early embryonic tissues in *CrebA* null embryos compared to WT. Only genes that are sufficiently expressed in at least one cell type in wild-type scRNA-seq are shown. The gray boxes correspond to genes whose expression in that tissue was too low in both genotypes to obtain a reliable fold change in gene expression.

Our new *CrebA* mutant stage 10 – 12 dataset contains all the cell types identified in the stage 10-12 *Drosophila* WT embryo atlas, with the single exception of apodemes (**Figure 3B, 2A**). Cells expressing all top three WT apodeme cell markers were few and mostly scattered throughout the *CrebA* mutant UMAP (**Figure S2C**), suggesting that we failed to recover this rare cell type in sufficient numbers in the early *CrebA* mutant dataset. Similarly, only one cell type identified in the early *CrebA* dataset was not present as a distinct cluster in the early wild-type sample – the optic lobe. Cells expressing the top three optic lobe marker genes were found within an epidermal cluster in the early wild-type dataset (**Figure S2D**), suggesting that these cells may be present and may require further sub-clustering to isolate. However, an optic lobe cluster was present in our late stage 13-16 WT dataset (**Figure S1A**). When we compare the average expression profiles from early WT cell types and early *CrebA* mutant cell types, the Spearman correlations between the average cell type expression were very high (0.96-0.99) across all cell types, even those with naturally high expression of *CrebA* (**Figure S3**). Finding nearly all the same cell types in WT and *CrebA* mutant embryo samples as well as the high correlation in average gene expression in pairwise comparisons of these cell types indicates that *CrebA* plays no role in the establishment of early cell fates.

To gain insight into the biological processes most impacted by *CrebA* loss, we found the differentially expressed genes between wild-type and *CrebA* mutants (**Table S2**) and performed gene-set enrichment analysis (GSEA) (**Table S3, S4**) comparing the highest *CrebA* expressing tissues (SG, amnioserosa, plasmatocytes, fat body, trachea, glia, and hindgut) between WT and *CrebA* mutant embryos. The top five *CrebA* expressing WT tissues were highly enriched for genes that function in the secretory pathway when compared to the same tissues in the *CrebA* mutants; indeed the top four categories from the GSEA analysis of the SG, amnioserosa, plasmatocytes, fat body, and trachea were terms such as Golgi vesicle transport, ER to Golgi vesicle-mediated transport, vesicle mediated transport, and retrograde vesicle-mediated transport (**Figure 3C**). GSEA analysis of the glia and hindgut, two tissues also expressing high *CrebA*, revealed secretory genes to not be as significantly enriched in WT versus *CrebA* nulls, perhaps because *CrebA* is only beginning to be expressed in these tissues in early embryos. To test this hypothesis, we examined data from stage 13-16 wild type and *CrebA* mutant scRNA-seq datasets (**Figure S4A**), albeit a stage for which we have only a single *CrebA* mutant sample instead of two. Nonetheless, GSEA analysis reveals that the gene sets most expressed in late-stage wild type relative to late stage *CrebA* mutant glia and hindgut are secretory pathway (**Figure S4B**). In contrast, GSEA of what is enriched in *CrebA* mutant versus WT tissues (functions CrebA might repress) revealed no connections to secretion and no biological processes affected across the different tissues, although some terms did occur in more than one tissue type: mRNA processing, ribosomal biogenesis, translation, and nucleosome organization (**Figure 3C**).

We next examined the expression changes in individual SPCGs and other CrebA targets in *CrebA* null versus WT tissues. The expression of SPCGs was generally reduced in *CrebA* mutant tissues with the most profound expression decreases in the SG, amnioserosa, plasmatocytes, fat body, trachea, glia, and hindgut (**Figure 3D**) – the tissues that have the highest levels of *CrebA* expression in WT embryos (**Figure 2B**). Confirming previous in situ and microarray data (Fox *et al*. 2010; Johnson *et al*. 2020), expression of *sage*, *Xbp1* and *TudorSN* also went down; *sage* levels decreased in only the SG (the one tissue where *sage* is expressed) and levels of the other two genes decreased in multiple CrebA expressing tissues (**Figure 3D**). Taken together, the relative fold change in expression of SPCGs, *Xbp1*, and *TudorSN* across tissues correlated with levels of *CrebA* expression seen in WT embryos (**Figure 2D, 3D**). In contrast, minimal or no changes in SPCG expression were observed in tissues such as CNS, gut endoderm, germ cells and circular visceral mesoderm, which do not express *CrebA* (**Figure 2D, 3D**). Likewise, when we performed GSEA, we do not see biological pathways associated with secretory functions to be enriched in wild-type CNS, gut endoderm, germ cells and circular visceral mesoderm relative to *CrebA* mutant tissues (**Figure S4D, S4E**). Altogether, these analyses reveal that the primary function of CrebA is to boost secretory capacity by upregulating secretory pathway component genes (SPCGs) in the embryonic tissues in which it is expressed.

### CrebA is sufficient to induce SPCG expression in tissues where it is not normally expressed

The findings presented here and in earlier studies reveal that CrebA expression correlates with and is required for SPCG expression in multiple tissues during embryonic development (Abrams and Andrew 2005; Fox *et al*. 2010; Johnson *et al*. 2020). Importantly, ectopic expression of UAS-*CrebA* in epidermal stripes using the *engrailed*-Gal4 (*en*-Gal4) driver results in epidermal stripes of expression of all 14 downstream target SPCGs that have been tested (*SrpRα*, *Sec61β*, *Spase25*, *p24.1*, *γCOP*, *Tbc1*, *Wbl*, *Sel*, *Der1*, *Osbp*, *Ufc1*, *CG5021*, *CG8230*, *lqfR*) (Fox *et al*. 2010; Johnson *et al*. 2020). Likewise, elevated expression of CrebA in the trachea and ventral midline cells using *breathless*-Gal4 (*btl*-Gal4) to drive UAS-CrebA resulted in expression of the five SPCGs that were tested in those tissues (Fox *et al*. 2010). These studies suggest that CrebA could be sufficient to drive SPCG expression in any tissue. To fully test this idea, we asked if ectopic expression of CrebA can drive ectopic expression of SPCGs specifically in tissues with <u>no</u> detectable levels of *CrebA* expression in early WT embryos. We used tissue-specific Gal4 driver lines to express UAS-*CrebA* in mesoderm and somatic muscle (*twi*-Gal4 or *Mef2-*Gal4), in the nervous system (*elav*-Gal4), or in germ cells (*nos*-Gal4) and visually assessed changes in gene expression for five SPCGs: *Spase12*, *KdelR*, *βCop*, *Sec61β* and *SrpRa* (**Figure 4**). First, we drove UAS-CrebA expression using the same en-Gal4 driver used previously. With all the SPCGs tested, we could drive their ectopic expression in *engrailed* stripes in earlier stage embryos (stage 9 – stage 13); however, ectopic expression of these same SPCGs in engrailed stripes was less intense/obvious at later stages (**Figure 4B**; **S5B**). UAS-CrebA expression using both the *mef2*-Gal4 and *twi*-Gal4 resulted in ectopic expression of all five SPCGs in all mesodermally-derived tissues in which we could detect CrebA expression both early and late, suggesting that ectopic expression of CrebA in both somatic and visceral mesoderm, tissues that do not normally express CrebA, is sufficient to drive SPCG expression (**Figure 4C,D**; **S5E,F**). Twi-Gal4 also drives expression in the midgut endoderm, where we also observed ectopic expression of the SPCGs (**Figure 4D;S5F**). Interestingly, we got variable results with ectopic CrebA expression using *elav*-Gal4 (nervous system) and *nos*-Gal4 (germ cell), depending on the *SPCG* gene examined. With one SPCG – *Spase12* – we observed ectopic expression in the CNS with *elav*-Gal4 and in late-stage germ cells with *nos*-Gal4 (**Figure 4E,F**). We also observed CNS expression of *SrpRα* and *Sec61β*, but not *KdelR* or *βcop*, when CrebA expression was driven with *elav*-Gal4 (**Figure 4E**; **S5I**). No ectopic expression of *SrpRα*, *βcop* or *KdelR* was observed in germ cells using *nos*-Gal4 to drive ectopic CrebA expression (**Figure 4F**; **S5J**). Thus, CrebA is sufficient to boost SPCG expression in multiple tissues, including in cell types with normally undetectable CrebA expression; CrebA cannot, however, drive expression of every SPCG in all stages or in all cell types (**Figure 4; Figure S5**).

**Figure 4.**
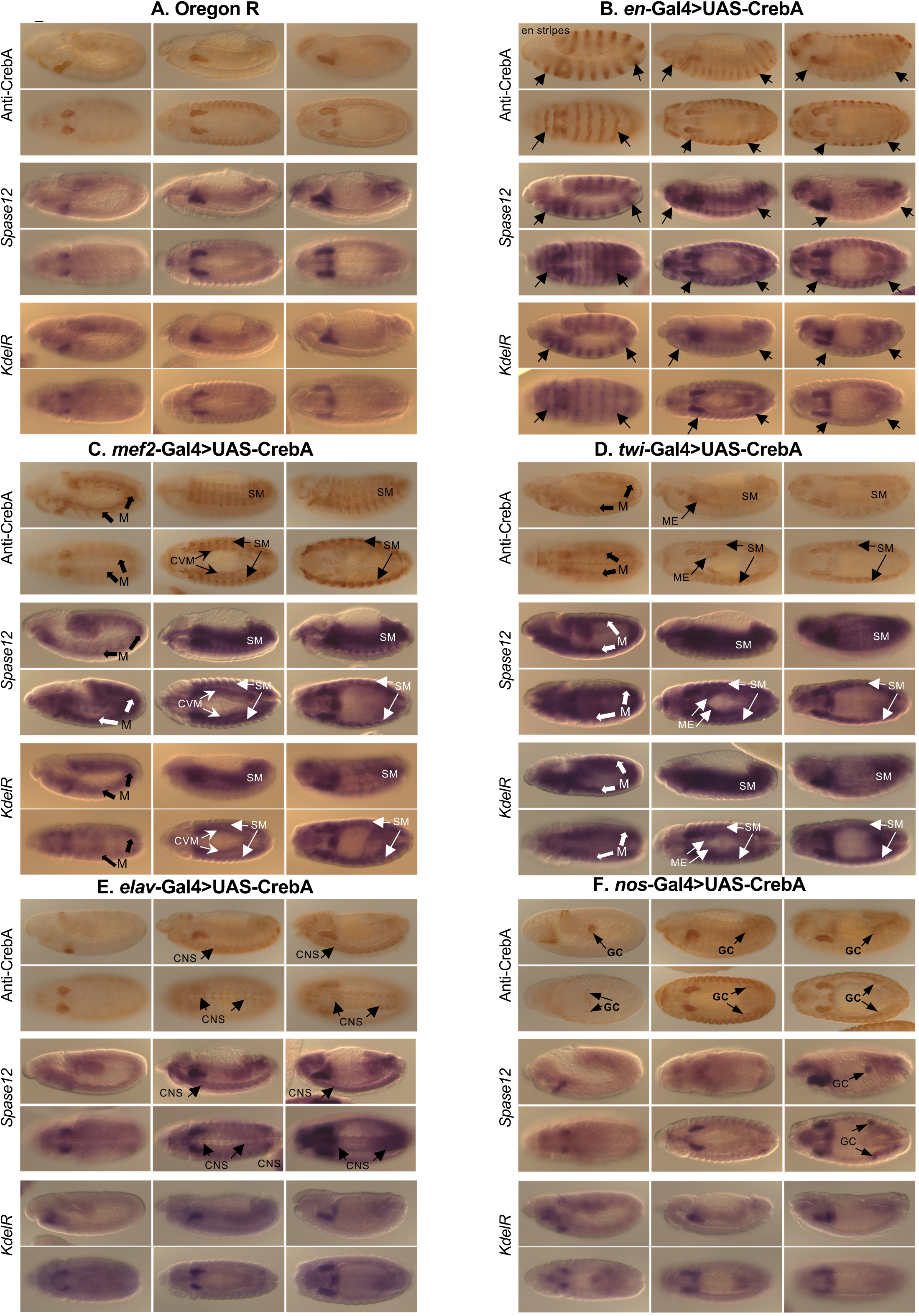
Ectopic expression of *CrebA* in tissues that do not normally express it can drive ectopic expression of target SPCGs. **A.** Oregon R (wild-type) embryos showing immunostaining for CrebA (row 1 – lateral views, 2 – ventral views) and mRNA accumulation of two SPCGs: *Spase 12* (row 3 – lateral views, 4 – ventral views) and *KdelR* (row 5 – lateral views, 6 -ventral views). **B.** *en*-Gal4>UAS-*CrebA* embryos showing immunostaining for CrebA (row 1 – lateral views, 2 – ventral views) and mRNA accumulation of two SPCGs: *Spase 12* (row 3 – lateral views, 4 – ventral views) and *KdelR* (row 5 – lateral views, 6 -ventral views). **C.** *mef2*-Gal4>UAS-*CrebA* embryos showing immunostaining for CrebA (row 1 – lateral views, 2 – ventral views) and mRNA accumulation of two SPCGs: *Spase 12* (row 3 – lateral views, 4 – ventral views) and *KdelR* (row 5 – lateral views, 6 -ventral views). **D.** *twi*-Gal4>UAS-*CrebA* embryos showing immunostaining for CrebA (row 1 – lateral views, 2 – ventral views) and mRNA accumulation of two SPCGs: *Spase 12* (row 3 – lateral views, 4 – ventral views) and *KdelR* (row 5 – lateral views, 6 -ventral views). **E.** *elav*-Gal4>UAS-*CrebA* embryos showing immunostaining for CrebA (row 1 – lateral views, 2 – ventral views) and mRNA accumulation of two SPCGs: *Spase 12* (row 3 – lateral views, 4 – ventral views) and *KdelR* (row 5 – lateral views, 6 -ventral views). **F.** *nos*-Gal4>UAS-*CrebA* embryos showing immunostaining for CrebA (row 1 – lateral views, 2 – ventral views) and mRNA accumulation of two SPCGs: *Spase 12* (row 3 – lateral views, 4 – ventral views) and *KdelR* (row 5 – lateral views, 6 -ventral views). Arrows highlight ectopic expression of CrebA and mRNAs. Abbrev. M = mesoderm, CVM = circular visceral mesoderm, S = somatic muscle, ME = midgut endoderm, CNS = central nervous system, GC = germ cells. For each set **A - F**, left panels are ∼stage 11, middle panels are ∼stage 13 and right panels are ∼stage 15.

### scRNA-seq captures tissue-specific expression changes in *CrebA* knockout

Multiple methods have been used to discover gene expression changes in *CrebA* null versus WT embryos as the tools and technologies have become available. Here, we compare our current scRNA-seq dataset, which enables the analysis of transcripts at single-cell resolution, with our datasets from whole embryo microarrays and whole mount in situ hybridization (Abrams and Andrew 2005; Fox *et al*. 2010; Johnson *et al*. 2020). We first compared the gene sets whose expression went down by scRNA-seq in any of the five tissues that express the highest levels of *CrebA* in the early embryo. We found that many genes went down in more than one CrebA-expressing tissue (**Figure 5A**). Of the genes that went down in the SG, 79.1% of those genes (125/158) also went down in at least one other tissue. Similar overlaps were seen with trachea (76.9%), amnioserosa (79.1%), and fat body (72.8%). Only 50.3% of the genes that went down in plasmatocytes also went down in at least one other tissue. This is largely because of the large number of genes that went down in expression in plasmatocytes relative to any other tissue. Similar levels of overlap were seen with genes that went up in *CrebA* mutants (**Figure 5B**). This indicates that most CrebA target genes are shared across the tissues that express high levels of CrebA protein.

**Figure 5.**
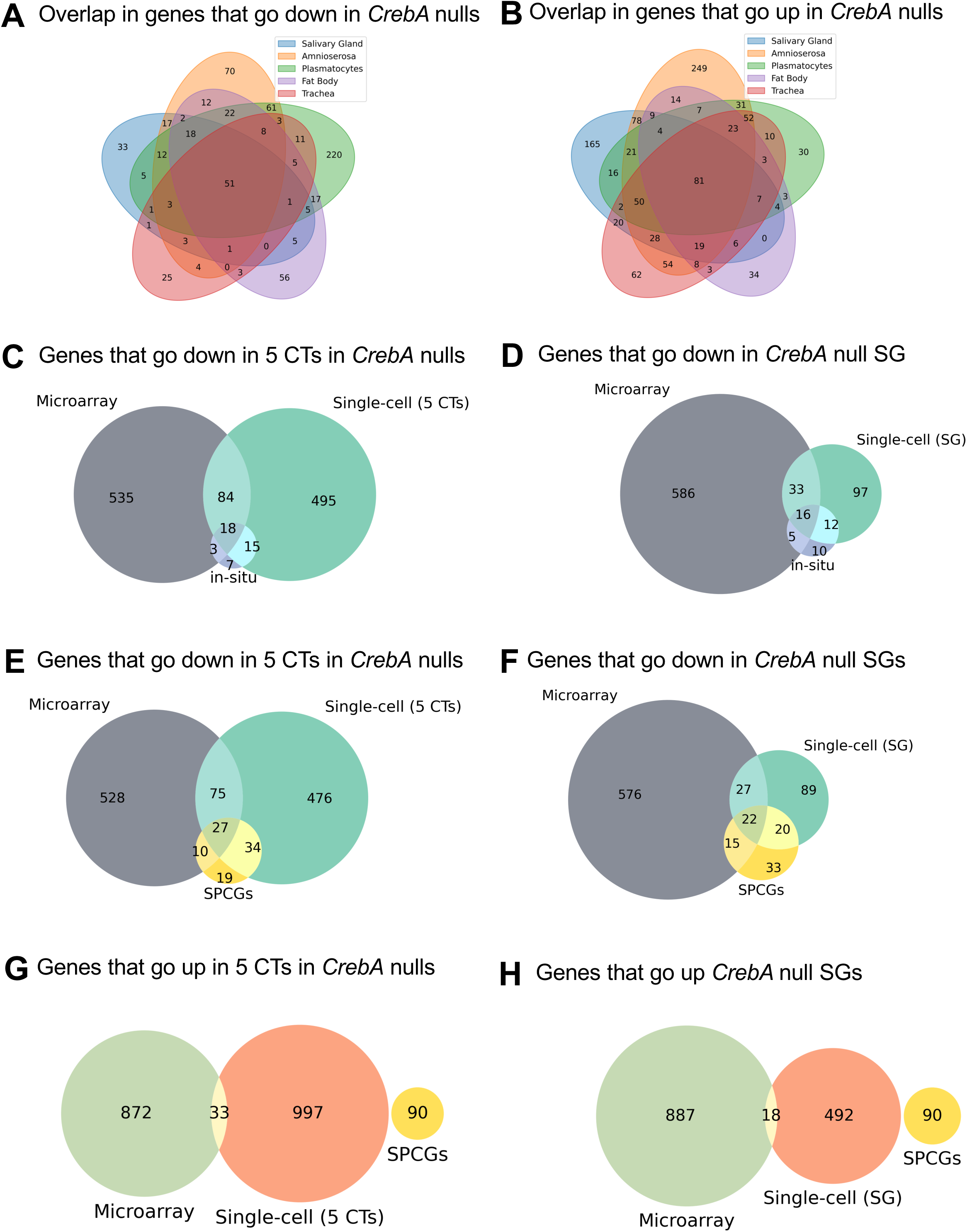
Comparisons of gene expression changes in the tissues expressing the highest levels of *CrebA* using different methodologies. **A.** Overlap in genes that go down in *CrebA* mutants relative to WT in the five early tissues expressing the highest levels of *CrebA* (SG, amnioserosa, plasmatocytes, fat body, trachea). **B.** Overlap in genes that go up in *CrebA* mutants relative to WT in the five early tissues expressing the highest levels of *CrebA*. **C.** Overlap in genes that go down in *CrebA* mutants relative to WT in the five early tissues expressing the highest levels of *CrebA* compared to genes that go down based on microarray analysis and on whole mount in situ hybridization analysis. **D.** Overlap in genes that go down in *CrebA* mutants relative to WT in the SGs compared microarray analysis and whole mount in situ hybridization analysis. **E.** Overlap in genes that go down in *CrebA* mutants relative to WT in the five early tissues expressing the highest levels of *CrebA* compared to microarray analysis of *CrebA* mutants and to the previously identified 90 SPCGs. **F.** Overlap in genes that go down in *CrebA* mutants relative to WT in the SGs compared to microarray analysis and to the previously identified 90 SPCGs. **G.** Overlap in genes that go up in *CrebA* mutants relative to WT in the five early tissues expressing the highest levels of *CrebA* compared to genes that go up based on microarray analysis and to the previously identified 90 SPCGs. **H.** Overlap in genes that go up in *CrebA* mutants relative to WT in the SGs compared to microarray analysis and to the previously identified 90 SPCGs.

We next examined the overlap in genes whose expression went down significantly in *CrebA* nulls based on scRNA-seq (logFC < -0.15 and p-val < 0.05) relative to genes whose expression went down based on whole embryo microarray (fold-change < -1.25 and p-val < 0.05). Overall, the overlap between the significantly down-regulated genes was smaller than we expected; only 16.7% of genes that went down in any of the five highest CrebA expressing tissues based on scRNA-seq also went down by microarray (**Figure 5C,E**). The overlap is better for the SG, where 31.0% of genes that went down in the SG by scRNA-seq also went down by whole embryo microarray (**Figure 5D,F**). When we compared scRNA-seq and microarray datasets with our in-situ dataset, we find that scRNA-seq analysis of the five highest CrebA expressing tissues shows higher overlap with CrebA-dependent genes identified by in situ (33/43 =76.7%) than microarray analysis does (21/43 = 48.8%) (**Figure 5C**). Likewise, scRNA-seq identified a greater number of SPCGs as going down in *CrebA* mutants (61/90 = 67.8%) than were identified by whole-embryo microarray analysis (37/90 = 41.1%; **Figure 5E**). The overlap between in situ data and SPCGs was also higher with scRNA seq data from the SG alone compared to the microarray data (**Figure 5D,F**). Of note, the overlap between scRNA-seq and microarray in terms of the genes that went up in *CrebA* mutants in either all five CrebA expressing tissues or in the SG alone was only 3.2 and 3.5% of genes (**Figure 5G,H**).

Many factors can explain why these methodologies detect different sets of CrebA targets. For example, the microarray experiments were from whole embryos collected over a larger time span (stages 10 – 16 embryos) (Fox *et al*. 2010); thus, we tested if more overlapping genes could be recovered if we included both the early and the late scRNA-seq datasets, again with the caveat that our late dataset came from only a single biological sample. When we did this for the SG only, we found that 29.9% of SG genes identified by early and late scRNA-seq were observed in the microarray data (**Figure S5**), whereas when we compared only the early scRNA-seq data, 31% of SG genes were also captured in the microarray. So, having a more comparable time span did not increase proportion of overlap. Both the microarray and scRNA-Seq approaches miss relevant targets found via a limited number of in situ experiments: microarrays because genes expressed in only a small subset of cells (e.g. SG) may be lost when transcript levels are averaged over the entire embryo, scRNA-seq because of transcript drop out or because of the limited depth of sequencing of individual cells, wherein rarer transcripts would be lost. Nonetheless, even with the more limited timed collections, scRNA-seq appears to recover more of the SPCGs and CrebA targets identified by in situ than were recovered by microarray in both SG only (47% and 65%, respectively) and in the combined five cell type data (77% and 68%, respectively), suggesting that the scRNA-seq is more sensitive than microarray for discovering CrebA downstream targets (**Figure 4**).

### CrebA binding and gene regulation in salivary gland cells

Previous ChIP-seq in vivo binding analyses of CrebA revealed that this TF binds many more genes than it regulates based on either microarray or in situ identified CrebA targets (Johnson *et al*. 2020). To further explore the relationship between CrebA binding and regulation of expression as well as to identify cis acting features that correlate with regulation, we integrated gene expression changes in the SG from scRNA-seq with ChIP-seq of CrebA SG-specific binding peaks. To identify the SG-specific binding regions, we found the ChIP-seq regions that overlap between sage-Gal>UAS-CrebA-GFP and fkh-Gal4>UAS-CrebA-GFP datasets and identified 5019 SG-specific CrebA binding regions in cells that express both *sage-*Gal4 and *fkh*-Gal4. We matched the SG-specific binding regions to the genes with their most 5’ transcription start site (TSS) closest to the center of the binding regions (both upstream and downstream and within 1kb).

Despite allowing each peak to potentially correspond to two genes, an overwhelming number of peaks are only assigned to either one or no genes (**Figure S7**). After matching peaks to genes, we found that CrebA binds to 2599 genes (**Figure 6A, 6B**). Among the 2599 CrebA bound genes (from the ChIP-seq), we identified 200 genes activated by CrebA and 176 genes repressed by CrebA (from the scRNA-seq) (**Figure 6A, 6B**). As described above, we found that scRNA-seq alone does not capture all genes that are activated by CrebA (**Figure 5D, F**), and different transcript capturing methods (microarray, in situs and scRNA-seq) capture genes missed by other methods. To maximize inclusivity of target genes activated by CrebA, we defined activated genes as those meeting any of the following criteria: 1) genes that show significant decrease in expression (logFC < -0.15 and p-value < 0.05) in *CrebA* mutant SG cells based on scRNA-seq, 2) genes previously identified as CrebA-activated via mRNA in situ hybridization experiments, or 3) genes that show a significant decrease in expression (fold-change < -1.25 and p-value < 0.05) in *CrebA* mutant embryos according to microarray, provided they have sufficient expression in the wild-type SG cells [>10 percent expression) and decrease in expression in CrebA mutant SG cells from scRNA-seq] (Abrams and Andrew 2005; Johnson *et al*. 2020). Due to extremely low overlap between genes that went up in *CrebA* knockout as identified by scRNA-seq and microarray (**Figure 5H**), we suspect many of the repressed genes from microarray are most likely not from SG cells, therefore we defined repressed genes as those that show significant increase in expression (logFC > 0.15 and p-value < 0.05) in *CrebA* mutant SG cells based on only scRNA-seq. This analysis revealed that only about 14% of genes that are bound are regulated by CrebA at early stages. These genes are labelled as CrebA bound and activated genes and CrebA bound and repressed genes, respectively, forming a direct downstream gene regulatory network for CrebA (**Table S5**). We observe that CrebA bound and activated genes encompass 73% of all activated genes whereas CrebA bound and repressed genes encompass only 35% of all repressed genes (**Figure 6A,B).** This suggests that CrebA is primarily a transcriptional activator in the SG. Interestingly, 76 out of the 90 SPCGs are directly bound by CrebA, with 54 of those showing reduced expression with loss of *CrebA*, further validating the importance of CrebA in directly upregulating the secretory machinery (**Figure 6A**). Also consistent with prior findings is that CrebA is a promoter proximal transcription factor that binds near the most 5’ TSS of the genes it regulates (**Figure 6C, 6D**).

**Figure 6.**
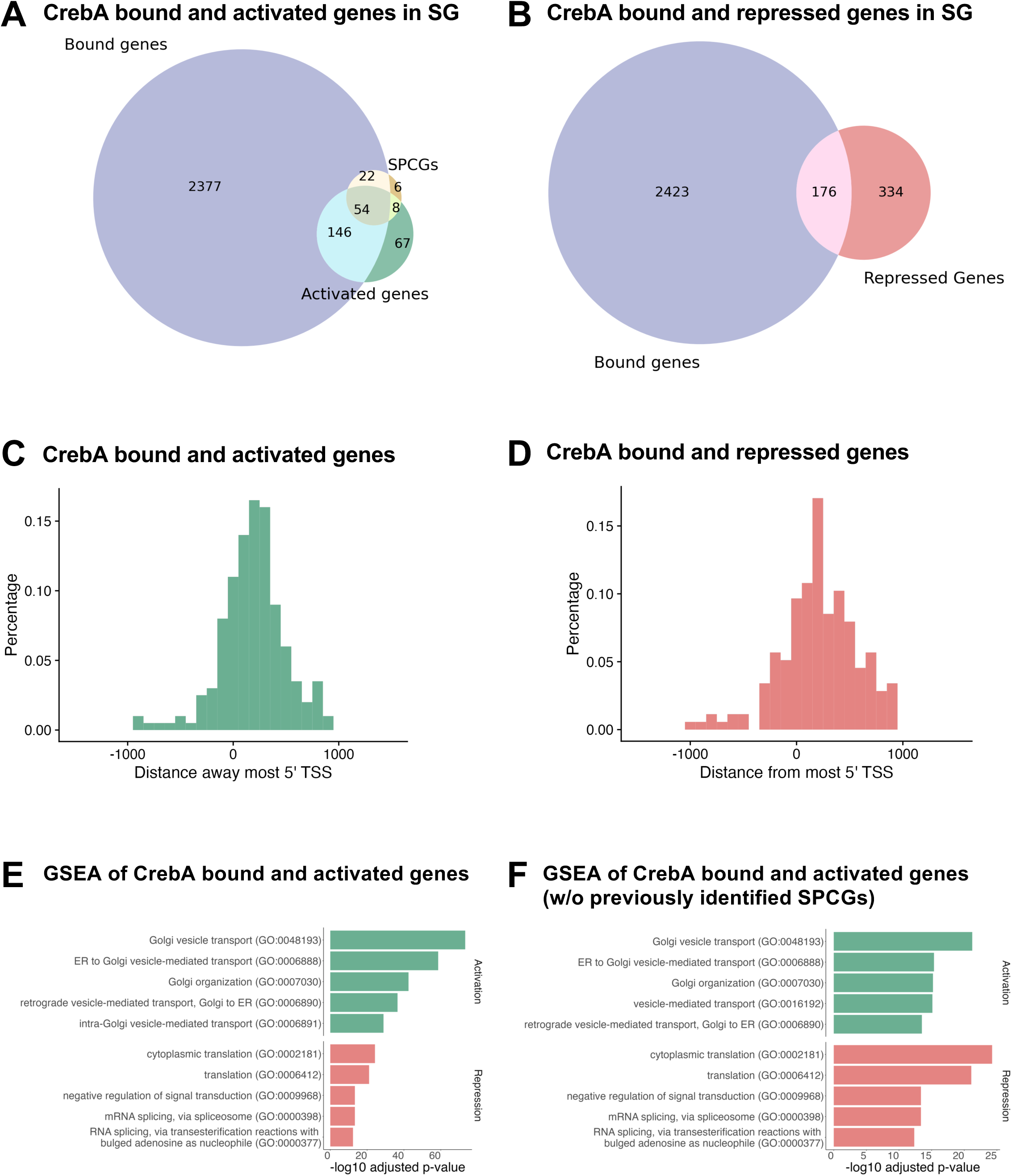
CrebA binding and changes in gene expression in *CrebA* mutants. **A.** Venn diagram showing the overlap in genes bound by CrebA in the SG and genes whose expression is reduced in the SG in *CrebA* mutants relative to WT and the 90 previously identified SPCGs. **B.** Venn diagram showing the overlap in genes that are bound by CrebA in the SG and genes whose expression goes up in the SG in *CrebA* mutants relative to WT. **C.** Histogram showing the distance between the CrebA binding peak to the 5’-most transcription start site (TSS) of genes whose expression is reduced in *CrebA* mutants relative to WT. **D.** Histogram showing the distance between the CrebA binding peak to the 5’-most TSS of genes whose expression is elevated in *CrebA* mutants relative to WT. **E.** GSEA of CrebA bound and activated genes and CrebA bound and repressed genes in SG. **F.** GSEA of CrebA bound and activated genes and CrebA bound and repressed genes after the removal of previously identified SPCGs.

To learn more about the classes of genes directly regulated by CrebA, we performed gene-set enrichment (GSEA) analysis using EnrichR (Chen *et al*. 2013), and found that CrebA bound and activated genes (based solely on SG scRNA-seq) are largely enriched in the secretory pathway whereas CrebA bound and repressed genes are largely enriched in translation and post-transcriptional modification (**Figure 6E**). This further supports the role of CrebA as a direct activator of secretory capacity. We also removed the CrebA bound and activated SPCGs (genes previously identified as SPCGs; Johnson et al., 2020) from the enrichment analysis, as they constituted a big portion of CrebA bound and activated genes. GSEA revealed that the remaining genes were still highly enriched in secretory pathway functions, suggesting that CrebA not only directly activates the secretory machinery components previously discovered but also other genes that contribute to secretory capacity (**Figure 6F**).

Next, we sought to identify features that distinguish CrebA bound and activated genes, CrebA bound and repressed genes, and CrebA bound and unchanged genes in the SG. We carried out DREME analysis of CrebA binding peaks (250 bp upstream and downstream of the center of the binding regions) for all the activated genes, repressed genes, as well the 150 genes whose expression changed the least in *CrebA* mutant SG (**Figure 7**). This analysis identifies enriched motifs in each group of genes. Perhaps not surprisingly, we identified a unique enriched sequence motif that correlates <u>only</u> with genes that went down in the SG of *CrebA* mutants. This motif matches the CrebA consensus binding motif previously identified from in vitro binding studies (Abrams and Andrew 2005; Noyes *et al*. 2008; Nitta *et al*. 2015) (**Figure S8A**). All other enriched motifs that emerged from these studies were not overly represented in any one class of genes (bound and activated, bound and repressed, and bound and unchanged), and included binding consensus motifs for E-box bHLH proteins (**Figure S8B**). The low enrichment of CrebA consensus binding motif from the bound and unchanged gene set or from the bound and repressed gene sets further supports our model that CrebA functions primarily as a transcriptional activator.

**Figure 7.**
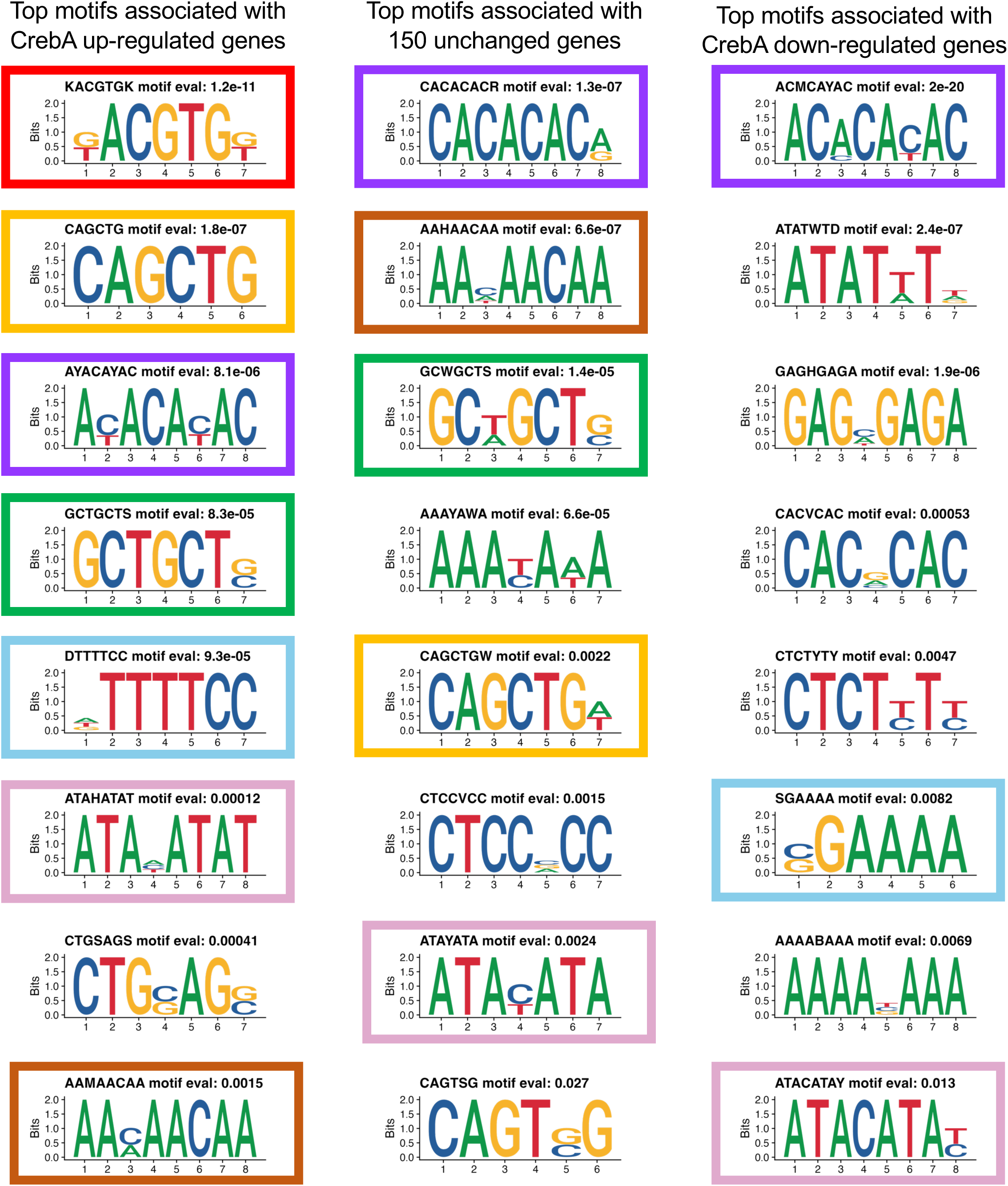
DREME analysis of different classes of CrebA bound genes in the SG. Left column: Conserved motifs identified by DREME analysis of CrebA peaks (+/- 250 bp) from the center of binding peaks of genes whose expression is reduced in the SG in *CrebA* mutants relative to WT. Middle column: Conserved motifs identified by DREME analysis of CrebA peaks (+/- 250 bp) from the center of binding peaks of 150 genes whose expression is least changed in the SG in *CrebA* mutants relative to WT. Right column: Conserved motifs identified by DREME analysis of CrebA peaks (+/- 250 bp) from the center of binding peaks of genes whose expression is elevated in the SG in *CrebA* mutants relative to WT. Boxed regions indicate motifs that are either unique to CrebA upregulated genes (red box) or are shared among the different regulatory classes of genes (yellow, purple, green, blue, pink boxes).s

## Discussion

### CrebA increases secretory capacity in all tissues in which it is highly expressed

In this study, we combined scRNA-seq with previously generated ChIP-seq and microarray data to identify embryonic tissues with high expression of CrebA and the secretory machinery, study the transcriptional effect of CrebA knockout at tissue-specific resolution, and identify direct downstream targets of CrebA in the SG and, likely, other embryonic tissues. Based on expression of SPCGs and *CrebA* in the previously published *Drosophila* embryo scRNA-seq atlas, we found that *CrebA* expression correlates with the expression of SPCGs across embryonic tissues. Furthermore, we identified several cell types that have high expression of *CrebA* and SPCGs that support their cellular functions. One of the highest expressing tissues is the SG, a dedicated secretory organ that will ultimately produce and secrete large quantities of the glue proteins that allow the mature larva/pupa to attach to a solid substrate during metamorphosis. Earlier, the SG produces secretions that allow for embryo hatching, protection against microbes, as well as the secreted enzymes and lubricants that aid in food digestion (Loganathan *et al*. 2021). Plasmatocytes, the embryonic immune cells that migrate throughout the embryo and secrete the basal ECM surrounding many embryonic organs (Ratheesh *et al*. 2015; Matsubayashi *et al*. 2017), are the earliest cells to express CrebA and have the greatest number of genes whose expression goes down with *CrebA* loss. Tracheal cells also express elevated levels of *CrebA*; these cells will secrete an apical chitinous extracellular matrix that serves to shape and regulate the expansion of tracheal branches (Tsarouhas *et al*. 2007; Hayashi and Kondo 2018). The fat body, which also expresses elevated levels of CrebA, not only functions as an endocrine organ supporting tissue growth (Okamoto *et al*. 2009), it has also been shown to synthesize and secrete at least one of the ECM proteins that ends up in the tracheal lumen (Dong *et al*. 2014). Although not considered a classic secretory organ, the amnioserosa secretes at least one key signaling molecule required for dorsal patterning (Lacy and Hutson 2016); perhaps this study suggests that additional secretory functions for the amnioserosa that await discovery. When CrebA is knocked-out, there are significant decreases in SPCG expression in all the high *CrebA* expressing cell types including the SG, trachea, plasmatocytes, fat body, and amnioserosa. Indeed, secretory functions are by far the most affected process in these tissue types with loss of CrebA. Non-*CrebA* expressing cell types such as CNS, gut endoderm, germ cells, circular visceral mesoderm and somatic muscle show no detectable change in overall SPCG expression in *CrebA* null embryos.

### CrebA functions as a transcriptional activator

Whereas the genes that require CrebA for expression are primarily linked to secretory capacity, an analysis of the genes whose expression increased in *CrebA* mutants from either previous microarray studies (Fox *et al*. 2010) or the current scRNA-seq analysis did not reveal any class of genes to be specifically enriched. Corresponding whole mount in situ experiments revealed that CrebA is required to achieve high levels of SPCG expression in many tissues (Abrams and Andrew 2005; Fox *et al*. 2010; Johnson *et al*. 2020) and that ectopic expression of CrebA is sufficient to activate target gene expression in cell types that do not normally express CrebA (Fox *et al*. 2010; Johnson *et al*. 2020) this study), supporting a role for CrebA as a transcriptional activator. Nonetheless, both microarray and scRNA-seq revealed that genes that go up in the *CrebA* mutants are even more numerous than the ones that go down. From the observation that CrebA binds to a much larger fraction of CrebA-activated genes than repressed genes and that the CrebA consensus binding motif identified from in vitro studies in the three different groups is only highly enriched in the binding regions of CrebA activated genes, we propose that CrebA primarily functions as a transcriptional activator. We further propose that the observed increases in gene expression observed in *CrebA* nulls are simply either a consequence of freeing up the general transcriptional machinery and/or because CrebA activates a transcriptional repressor.

Previous in situ hybridization studies showed that expression of every tested SPCG goes down in the SG and other tissues in *CrebA* null embryos, indicating that CrebA is essential to boost SPCG expression (45 SPCGs tested; see (Johnson *et al*. 2020)). Moreover, ectopic expression of *CrebA* in engrailed stripes is sufficient to drive striped expression of every SPCG previously tested (14 SPCGs); we find this is also true for the new SPCGs tested using the *en*-Gal4 driver in this study. Here, we demonstrate that ectopic CrebA expression in mesoderm and mesodermally-derived tissues (using *twi*- and *mef2*-Gal4) also results in ectopic expression of all the SPCGs we tested, with expression in the CVM and somatic muscle – tissues that do not express detectable levels of *CrebA* in WT. This suggests that CrebA is sufficient for SPCG activation in the context of any Drosophila embryonic cell type. However, with CrebA expression in the CNS using *elav*-Gal4, only three of the five SPCGs we tested had ectopic CNS expression. Likewise, CrebA expression in the GCs using *nos*-Gal4 resulted in GC SPCG expression of only one of the tested SPCGs and their GC expression was observed in only late-stage embryos. Our interpretation of these findings is that CrebA is sufficient to bind and activate target gene expression as long as the chromatin remains accessible – explaining the ectopic expression observed in the CNS with only a subset of SPCGs – or as long as other more general transcription blocking mechanisms are not activated, as we believe could be occurring in the early GCs. Indeed, GC transcription is known to be blocked by several mechanisms in the GC primordia of Drosophila and other higher organisms (Nakamura and Seydoux 2008).

### CrebA/Creb3L proteins are potent activators of secretory capacity across species, but they may not be the only activators

Since the original discovery that CrebA upregulates secretory capacity in the Drosophila SG by upregulating expression of SPCGs (Abrams and Andrew 2005), numerous studies in higher organisms have implicated the CrebA orthologues (known as the Creb3L family of bZip TFs) in boosting secretory capacity across a broad range of tissues from bone and cartilage to the liver and pituitary glands (Murakami *et al*. 2009; Saito *et al*. 2009; Barbosa *et al*. 2013; Reiling *et al*. 2013; Khetchoumian *et al*. 2019; Sampieri *et al*. 2019). Interestingly, a recent study in zebrafish revealed that another related TF, known as Xbp1, can also do the same job (Wang *et al*. 2024). This study showed that whereas Creb3L1 and Creb3L2 play largely overlapping roles in boosting SPCG expression in the notochord, Xbp1 performs this function in the hatching gland. Like the mammalian Creb3L proteins, Xbp1 was initially identified as a mediator of the unfolded protein response (UPR); although it too has been implicated in secretory machinery biogenesis (Lee *et al*. 2005). We find that in fly embryos, CrebA is the major regulator of SPCG expression, and its expression correlates with that of SPCGs in almost every tissue (this study; (Abrams and Andrew 2005). In flies, Xbp1 is a direct downstream target of CrebA in the SG and several other embryonic tissues (Johnson *et al*. 2020). Correspondingly, whereas CrebA loss has major consequences on secretory function all tissues that have been analyzed, loss of *Xbp1* has only minor consequences (Abrams and Andrew 2005; Johnson *et al*. 2020). Nonetheless, an exception to CrebA regulation of SPCG/secretory function in fly embryos could be in the garland cells (specialized renal cells for filtering the hemolymph); in later stage embryos, these cells have very high levels of *SPCG* expression, but low levels of *CrebA* (**Figure S1**). Notably, garland cells have the highest levels of *Xbp1*. So perhaps secretory capacity is also regulated by either CrebA or Xbp1 in different tissues in the fly. It is worth noting that CrebA and Xbp1 share the same consensus binding motif and both have bZIP DNA binding domains, suggesting similar targets and similar mechanisms of gene regulation.

### Limitations to this study

We note that there are limitations to our investigations. In this study, we reconstructed the direct functional network downstream of CrebA in the SG through the integration of scRNA-seq and SG specific CrebA ChIP-seq. However, we also acknowledge that SG is not the only embryonic cell type with high expression of CrebA, there are also amnioserosa, plasmatocytes, FB, and trachea. To generate the direct CrebA transcriptional network for the other cell types, we would need to generate cell-type specific CrebA binding peaks, using other tissue specific Gal4 lines to drive expression of GFP-tagged UAS-CrebA, as we have done for SG. In addition, our pipeline for calling CrebA binding even in the SG is not perfect. There are eight SCPGs that were previously found to be bound by CrebA via manual inspection (Johnson *et al*. 2020) but are not found to be bound via our pipeline (Arf79F, Pdi, Sar1, Sec23, Spase22-23, Srp72, TBC1D23, p24-2). Some of the main reasons for the discrepancy include no overlapping regions between *fkh*-Gal4 and *sage*-Gal4 driven UAS-*CrebA*-GFP expression experiments, the distance between TSS and the center of overlapping binding region is slightly above 1kb and another neighboring gene with TSS that is closer to the center of overlapping binding region.

We also note that we do not fully understand what molecular features distinguish a directly bound and activated CrebA target gene from a gene that is simply bound by CrebA. Almost certainly, CrebA binding is linked to accessible chromatin and the presence of a CrebA consensus or near consensus motif. Indeed, we observe specific enrichment of the CrebA consensus binding motif in promoter proximal regions in the set of genes that go down in *CrebA* mutant embryos; these sites also occur, however, in a smaller subset of bound genes whose expression either doesn’t change or goes up. We do note that among the motifs also present in CrebA bound regions is an EBox-bHLH consensus motif. In the SG, this motif is likely linked to Sage, a bHLH TF expressed specifically and to high levels in the SG, and whose expression goes down in the SGs of *CrebA* mutants. Sage activity could create the open chromatin states associated with non-functional CrebA binding and could contribute to the reduced expression of genes not directly bound by CrebA. Interestingly, CrebA appears to be sufficient to activate target gene expression even in cell types that do not normally express detectable CrebA, suggesting that if other TFs are required for CrebA driven expression, these TFs are either broadly expressed or activated by CrebA.

In summary, we have generated a novel scRNA-seq dataset for *CrebA* mutant embryos primarily in stage 10-12 and investigated the transcriptional consequences of *CrebA* knocked-out in embryonic tissues. This data provides the community a valuable resource to further explore the consequences of CrebA knock-out in various cell types. Integrating the genes that were significantly changed when CrebA is knocked-out with SG specific CrebA binding data, we inferred the direct downstream network of CrebA in SG during embryonic stage 10-12. This result further validates the importance of CrebA as a direct activator of secretory function in the SG and in other organs with high secretory activity.

## Acknowledgments

We thank Katherine Hutchings, JiHoon Kim, and Vidya Ajay for critical reading of the manuscript and for helpful suggestions during multiple stages of the project. We thank Nathaniel Laughner for converting the old gene names from the CrebA microarray studies to the newest Flybase ID/gene names and for his suggestions on the project. We thank Flybase for the gene annotations and current versions of the genome that were used for all of our data analysis (Flybase 1995). We thank the Drosophila Genomics Resource Center (DGRC) for the cDNA clones used in this study. We gratefully acknowledge support for this work from NIH RO1 DE013899 to D.J.A. and a Johns Hopkins Cell Biology Award to D.J.A. The authors declare no competing financial interests.

## Data availability

All raw data and processed data will be made available through GEO. Analysis scripts and processed data will be made available on Github.

## Supplemental Figure Legends

**Figure S1. Analysis of *CrebA* and SPCG expression across tissues from late-stage (stage 13-16) WT single cell RNA sequencing (scRNA-seq). A.** UMAP showing that all the major cell types are represented in the data (Peng et al., 2024). **B.** Violin plot of CrebA expression across tissues: tissues expressing highest levels are on the left, lowest on the right. **C.** In situ hybridization of *CrebA* (left panels) and CrebA immunostaining (right panels) at the stages captured by the late scRNA-seq analysis. The first and third rows are lateral views, the second and fourth are dorsal-ventral views. Abbreviations: SG – salivary gland, Tr – trachea, Es – esophagus, PV – proventriculus, Ph – pharynx, HG – hindgut, GL – glia, FB – fat body, St – stage. **D.** Late embryonic expression levels of *CrebA*, most known SPCGs, and three other known CrebA targets across late stage embryonic tissues.

**Figure S2. Analysis of CrebA mutant scRNA-seq replicates and of the cell types captured in distinct clusters. A.** UMAP showing the overlay of cells in *CrebA* mutant samples from biological replicas 1 and 2 after harmonization. **B.** UMAP from the merged *CrebA* mutant data showing the 40 identified clusters. **C.** UMAPs of *CrebA* mutant scRNA-seq data showing which cells express top apodeme markers (*CG7296*, *TwdIL*, *TwdlB*). **D.** UMAPs of WT scRNA-seq data showing which cells express the top optic lobe markers (*E(spl)m5-HLH*, *SoxN*, *Obp99a*).

**Figure S3. Gene expression profiles across different tissues do not change much in *CrebA* null versus WT early embryos.** Graph showing the Spearman correlation in gene expression profiles across all tissues in WT and *CrebA* mutant scRNA-seq data plotted against relative levels of *CrebA* RNA expression in different tissues.

**Figure S4. Analysis of gene expression changes in *CrebA* null versus WT in late embryos and in tissues that do not express *CrebA*. A.** UMAP from late stage *CrebA* null embryonic scRNA-seq showing that all major cell types found in the data. **B.** Gene set enrichment analysis (GSEA) reveals gene set categories enriched in the WT relative to *CrebA* null glia and hindgut at late stages (stage 13-16). **C.** Gene set enrichment analysis (GSEA) reveal gene set categories enriched in the glia and hindgut in *CrebA* null tissues relative to WT at late stages. **D.** GSEA analysis of WT versus *CrebA* null early gene expression in tissues that express the lowest levels of *CrebA* at early stages (stage 10-12). **E.** GSEA analysis of *CrebA* null versus WT early gene expression in tissues that express the lowest levels of *CrebA* at early stages.

**Figure S5 (two pages). βGal and SPCG expression in embryos using the different Gal4 lines to drive either *lacZ* or *CrebA* expression in the embryo. A.** Expression of three SPCGs in Oregon R embryos. **B.** Expression of the same three SPCGs in embryos using the *en*-Gal4 driver to drive UAS-*CrebA* expression in the embryo. **C.** Expression of βgal using the *mef2*-Gal4 driver to drive UAS-*lacZ* expression in the embryos. **D.** Expression of βgal using the *twi*-Gal4 driver to drive UAS-*lacZ* expression in the embryo. **E.** Expression of three SPCGs in embryos using the *mef2*-Gal4 driver to drive UAS-*CrebA* expression in the embryos. **F.** Expression of three SPCGs in embryos using the *twi*-Gal4 driver to drive UAS-*CrebA* expression in the embryos. **G.** Expression of βgal (top two rows) using the *elav*-Gal4 driver to drive UAS-*lacZ* expression in the embryos. Note that βgal is expressed in both the CNS and PNS, although we did not detect CrebA expression in the PNS using this same driver (see Figure 4E). **H.** Expression of βgal using the *nos*-Gal4 driver to drive UAS-*lacZ* expression in the embryo. Note that although βgal expression shows high level background staining with this driver, we did not detect high CrebA background staining with this driver (see Figure 4F). **I.** Expression of three SPCGs in embryos using the *elav*-Gal4 driver to drive UAS-*CrebA* expression in the embryos. **F.** Expression of two SPCGs in embryos using the *nos*-Gal4 driver to drive UAS-*CrebA* expression in the embryos. Top rows of embryos for each antibody or probe are lateral views. Bottom rows of embryos for each antibody or probe are mostly ventral views. The left most column in all sets are st 11 – 12, the middle column in all sets are stage 13. The right most column in all sets are ∼ st 15. Arrows highlight ectopic expression of lacZ/βGal and mRNAs. Abbrev. M = mesoderm, CVM = circular visceral mesoderm, S = somatic muscle, ME = midgut endoderm, CNS = central nervous system, GC = germ cells. For each set **A - F**, left panels are ∼stage 11, middle panels are ∼stage 13 and right panels are ∼stage 15.

**Figure S6. Overlap of genes that go down in *CrebA* nulls found using microarray and genes that go down in *CrebA* null SGs (both early and late stages) with scRNA-seq and with SPCGs.**

**Figure S7. Number of CrebA binding peaks observed with 0, 1 and 2 nearest genes whose 5’ ends map within 1 kb.** Note that a significant number of CrebA peaks have no nearby genes and only a small fraction have two nearby candidate genes.

**Figure S8. Enriched motifs in genes bound and whose expression goes down with loss of *CrebA* and the SG-expressed TFs that are known to bind each motif. A.** The top enriched motif found in CrebA bound genes that go down in *CrebA* null embryos matches the consensus motif for TFs expressed in the SG (green bars). Three of these (CrebA and Max) have enriched SG expression (dark green). **B.** The second most enriched motif found in genes whose expression is reduced in *CrebA* mutants matches the consensus motif for TFs expressed in the SG (green bars). Sage has enriched SG expression (dark green).

## Supplemental Figure Legends

**Table S1. Marker genes for the clusters in stage 10-12 *CrebA* mutant scRNA-seq atlas.**

**Table S2. Differentially expressed genes between stage 10-12 wild-type and *CrebA* mutant cell types.**

**Table S3. GSEA results for stage 10-12 wild-type cell types.**

**Table S4. GSEA results for stage 10-12 *CrebA* mutant cell types.**

**Table S5. Downstream targets of CrebA**

